# Disequilibrium in Gender Ratios among Authors who Contributed Equally

**DOI:** 10.1101/241554

**Authors:** Nichole A. Broderick, Arturo Casadevall

**Affiliations:** Department of Molecular & Cell Biology, University of Connecticut, Storrs, CT; Department of Molecular Microbiology and Immunology, Johns Hopkins School of Public Health, Baltimore, MD

**Author notes:** NAB and AC contributed equally. Author order was determined both alphabetically and in order of increasing seniority. Address correspondence to either: Nichole A. Broderick, University of Connecticut, Department of Molecular & Cell Biology, 91 North Eagleville Road, Unit 3125, Biology/Physics Building 304 Storrs, CT 06269-3125; Arturo Casadevall Johns Hopkins School of Public Health, Department of Molecular Microbiology and Immunology, 615 N. Wolfe Street, Room E5132, Baltimore, Maryland 21205.

**Keywords:** gender, publication, authorship, biomedical, research

## Abstract

In recent decades, the biomedical literature has witnessed an increasing number of authors per article together with a concomitant increase of authors claiming to have contributed equally. In this study, we analyzed over 3000 publications from 1995–2017 claiming equal contributions for authors sharing the first author position for author number, gender, and gender position. The frequency of dual pairings contributing equally was male-male > mixed gender > female-female. For mixed gender pairs males were more often at the first position although the disparity has lessened in the past decade. Among author associations claiming equal contribution and containing three or more individuals, males predominated in both the first position and number of gender exclusive groupings. Our results show a disequilibrium in gender ratios among authors who contributed equally from expected ratios had the ordering been done randomly or alphabetical. Given the importance of the first author position in assigning credit for a publication, the finding of fewer than expected females in associations involving shared contributions raises concerns about women not receiving their fair share of expected credit. The results suggest a need for journals to request clarity on the method used to decide author order among individuals claiming to have made equal contributions to a scientific publication.

---

In recent decades, the number of authors per publication has increased steadily [1; 2]. The causes for these trends include the higher production of scientific information by research teams [3] and increase in the data content of published papers [4; 5], which in turn usually requires contributions by additional scientists. This increase in number of authors per article has raised questions about credit allocation. Author order in an article byline is the major mechanism for assigning credit when there are more than one author. In general, the first author is the individual who has done most of the work and that individual traditionally receives most credit for publication. This in turn has resulted in an increase of authors claiming equal credit in author byline positions, which has posed vexing questions as to how credit should be apportioned [6; 7]. Analysis of articles in 5 medical journals showed that whereas papers listing equal contributions comprised less than 1% of publications in 2000, that by 2009 this trend had increased to 3.6–8.6% depending on the journal [8]. For the journal Gastroenterology 21% of papers from 2011–2102 indicated two or more authors contributing equally [9]. Hence, statements of equal contribution by more than one author is an increasingly common mechanism for sharing credit as the size of research teams increase in the biomedical sciences.

The shared authorship phenomenon is an important issue to study because the ability of junior investigators to publish first author papers is usually a necessary step for securing positions, funding, and receiving credit. To date very little scholarly work has been done to understand the mechanisms used in sharing credit allocation. In particular, we were interested in trends involving the sharing of equal contributions among authors differing in gender, since inequities in distribution could translate into differences in gender recognition for scientific accomplishment. Numerous studies have documented underrepresentation of women in academic faculty and in scientific positions, especially at the more senior ranks [10; 11; 12; 13]. Although the mechanisms for these trends are complex one possibility is that they receive less credit for their scientific work [12]. Several studies have documented gender differences in the frequency of first authors, with women less likely to occupy the first position [14; 15]. A large study of Swedish scientists revealed that women are more likely to be middle authors and less likely to be senior authors [16]. Hence, the available evidence suggests that disparities exist in gender contribution and position to the author byline of scientific publications.

In this study, we analyzed the gender order of publications where two or more individuals shared the first author position by stating that they had contributed equally. The expectation from equal contribution is that the order of author gender will be equally distributed or perhaps follow some ordering convention such as alphabetical order. Instead, we found a predominance of males at the first author position irrespective of whether first authorship was shared by two or more scientists. Furthermore, male-male pairings and all male chairing of equal credit was far more frequent that corresponding female associations. The finding of disequilibrium in gender ratios among authors who contributed equally suggests that inequities in credit sharing may be a contributing factor to the continuing gender imbalances reported for academic positions, grant funding and awards. The results suggest a need for more clarity and transparency in stating how author position is selected when more than one author share equal credit.

## Materials and Methods

The study was done in three stages. First, we undertook a cursory review of publications using the Google Scholar search engine with the keywords ‘contributed equally’ to familiarize ourselves with the variables involved and get a sense as to whether there were differences in how often males and females shared first author positions. This stage involved analyzing several hundred publications, which identified 57 publications that had one or more co-first authors (listed in Table 1 as results from ‘early searches’). This initial analysis revealed that whereas our initial interest was in gender positions among mixed gender pairs contributing equally there were many publications with more than two authors, suggesting the need for analyzing different journals. In the second stage, we undertook a systematic search of papers using two search strategies. One strategy used Google Scholar to search for the keywords ‘contributed equally’ and a specific journal name. The second strategy searched for the phrase ‘contributed equally’ in individual journal websites. After finding several hundred publications, we compared the results of the two search strategies and found discrepancies. Specifically, the Google Scholar search strategy was returning a higher frequency of male-female (M:F) orderings among those chairing first position than the in journal website search strategy. Inspection of the identified articles revealed that that the Google Scholar search strategy was returning more older papers suggesting a temporal variable to male-female author orderings, a finding that was subsequently confirmed at the conclusion of the study. The third stage of the study involved adding more papers using both the Google Scholar search and journal website strategies with the searches targeting specific years for those years where few papers were initially identified.

**Table 1.** Summary of data on authors listed a contributing equally.

| Journal Title | Article Statistics |  |  | Contributed equally = 2 |  |  |  | Contributed equally > 2 |  |  |  |
| --- | --- | --- | --- | --- | --- | --- | --- | --- | --- | --- | --- |
|  | Total | Unknown | Usable | mm | mf | ff | fm | m first | f first | all m | all f |
| Biophysical J | 101 | 2 | 99 | 60 | 17 | 5 | 9 | 5 | 0 | 3 | 0 |
| Cell Reports | 105 | 1 | 104 | 41 | 12 | 11 | 15 | 12 | 9 | 3 | 1 |
| Curr Biol | 103 | 4 | 99 | 41 | 23 | 14 | 12 | 2 | 4 | 2 | 1 |
| eLife | 90 | 2 | 88 | 25 | 18 | 15 | 11 | 5 | 6 | 8 | 0 |
| J Biol Chem | 300 | 28 | 272 | 99 | 52 | 35 | 48 | 15 | 11 | 6 | 6 |
| J Cell Bio | 101 | 3 | 98 | 22 | 35 | 14 | 17 | 4 | 4 | 2 | 0 |
| J Clin Invest | 121 | 6 | 115 | 42 | 19 | 11 | 19 | 12 | 6 | 6 | 0 |
| J Exp Med | 210 | 13 | 197 | 65 | 39 | 26 | 30 | 11 | 8 | 11 | 7 |
| J Immunol | 308 | 10 | 298 | 89 | 76 | 59 | 48 | 16 | 3 | 5 | 2 |
| mBio | 100 | 2 | 98 | 27 | 25 | 12 | 15 | 8 | 7 | 3 | 1 |
| Nature | 104 | 6 | 98 | 44 | 12 | 7 | 8 | 14 | 3 | 10 | 0 |
| PLOS Bio | 110 | 9 | 101 | 39 | 18 | 13 | 17 | 6 | 5 | 3 | 0 |
| PLOS Comp Bio | 95 | 2 | 93 | 45 | 14 | 6 | 11 | 3 | 4 | 10 | 0 |
| PLOS Genet | 186 | 8 | 178 | 52 | 26 | 16 | 23 | 20 | 35 | 6 | 0 |
| PLOS Negl Trop Dis | 105 | 6 | 99 | 32 | 13 | 21 | 19 | 8 | 4 | 2 | 0 |
| PLOS Pathogen | 179 | 9 | 170 | 35 | 39 | 33 | 25 | 18 | 9 | 7 | 4 |
| PNAS | 411 | 14 | 397 | 151 | 66 | 44 | 61 | 30 | 19 | 22 | 4 |
| Science | 128 | 11 | 117 | 34 | 15 | 21 | 25 | 7 | 9 | 6 | 0 |
| Initial search<sup>1</sup> | 57 | 0 | 57 | 0 | 35 | 0 | 22 | 0 | 0 | 0 | 0 |
| Misc<sup>2</sup> | 120 | 2 | 118 | 57 | 27 | 14 | 12 | 4 | 2 | 1 | 1 |
|  | **3034** | **138** | **2896** | **1000** | **581** | **377** | **447** | **200** | **148** | **116** | **27** |

1<sup>1</sup>These set papers are from the early searches used to identify the variables in this study and only MF and FM numbers were recorded.
2<sup>2</sup>Miscellaneous includes a variety of journals for which only a few articles were obtained. These journals with the number of articles in parenthesis are: American Journal of Pathology (1), Angewandte Chemie (17), Biochemical and Biophysical Research Communications (2), Blood (1), BMC Bioinformatics (1), BMC Proceedings (1), BMC Systems Biology (1), Brain Pathology (1), Cancer Research (2), Cell (6), EMBO Journal (4), European Journal of Immunology (1), FEBS Letters (27), Genes and Development (5), Genome Research (1), Hepatology (1), International Journal of Cancer (2), Journal of Bone and Mineral Research (2), Journal of Cell Science (2), Journal of Molecular Biology (1), Journal of Physiology (2), Memórias do Instituto Oswaldo Cruz (15), Nature Biotechnology (1), Nature Cell Biology (1), Nature Genetics (15), Nature Materials (1), Nature Medicine (4), PLOS One (1), Protein Science (1).

One of the two coauthors inspected each article manually. Determination of author gender was done by searching for images of an individual’s name using the Google search engine, which was adequate to assign gender for 97% of papers examined. Searches were narrowed by including the name of the research institution or research subject among the search words. In many instances, we were able to locate the individual by finding the website for the laboratory producing the paper. For those individuals whose gender could not be identified the major cause for failing identification was the absence of a photograph on the web. For each paper, we recorded the country of origin based on the country of the corresponding author, the gender of individuals sharing the first authorship, the year of publication, and whether the order of authors sharing equal credit was alphabetical. We estimate that the analysis of each entry averaged approximately 5 minutes, since each publication needed manual inspection to confirm that the search engine was correct and this often necessitated inspecting the PDF version of the publication, as author contributor information was not often available for the online format versions.

At the beginning of this study, the Pubmed database had approximately 25,000,000 million entries, which required a sampling size of 2400 articles to achieve a confidence interval of 2 with a confidence level of 95% (https://www.surveysystem.com/sscalc.htm). At the completion of the study, we had analyzed 2897 usable articles, which gave us a confidence interval of 1.82 at the 95% confidence level. However, given that only 0.8% of all papers in PubMed have first authors that contribute equally [9], this would reduce the size of the database of interest to approximately 200,000 publications. Hence, our database of 2897 usable publications represents about 1.45% of the available publications, which gives us a confidence interval of 2.38 at the 99% confidence level. The distribution of usable articles per year is shown in Supplemental Figure 1. Chi-square analysis used the on-line calculator http://www.socscistatistics.com/tests/chisquare/Default2.aspx

## Results

We analyzed 3035 scientific publications from 1995 to 2017 where two or more authors stated to have contributed equally. From this set, we identified the gender for each of the authors listed as contributing equally in 2897 publications, which provided our usable dataset (Table 1). Two authors were listed as contributing equally in 2406 (83%) publications while 491 (17%) listed three or more (Table 1).

For the 2330 publications where two authors contributed equally the most common gender pairing involved mixed gender (1028, 42.7%), which was closely followed by male-male pairings (1000, 46.6%). Female only author pairings were least frequent (447, 18.6%). Among the 1028 mixed gender pairings the male was listed first in 56.5% of publications. Assuming that individuals of both genders contributed equally and that this order was random, one would have expected roughly equal male-female and female-male pairings. Comparing the expected and observed gender ratios yielded a Chi-square statistic of p <0.001. Dividing these publications into those published 1995–2006 and 2007–2017 yielded 190 and 356 pairings for male-female order, respectively, and 103 and 322 pairings with female-male order, respectively. When these ratios were analyzed with the Chi-square statistic the difference between observed and expected ratios was significant in the 1996–2006 group (p < 0.001) but not in the 2007–2017 group (p = 0.8). Hence, male-female pairings were more common than female-male pairings and the frequency of these pairings deviated significantly from expected ratios if the assignments of author order were random, but the effect came primarily from publications in the 1996–2006 period. A plot of the frequency of the various author associations as a function of time provides a visual representation for this effect and for the changes as function of time (Figure 1).

**Figure 1.**
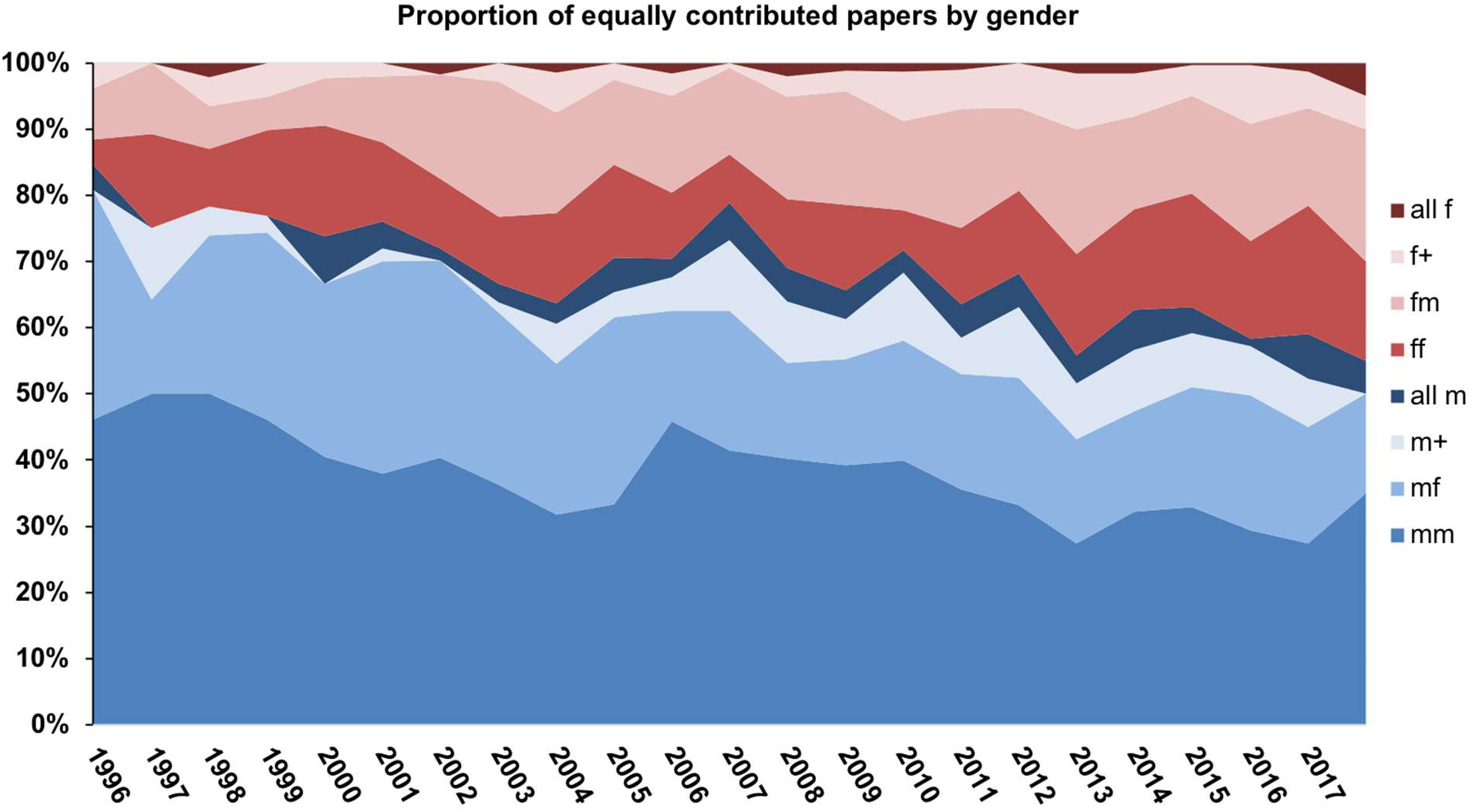
Proportion of equally contributed papers for the eight associations. The ‘fm’, ‘ff’, ‘mf’ and ‘mm’ categories represent pairings of female-male, female-female, male-female and male-male author orderings, respectively. The ‘all f’ and ‘all m’ represent groupings of more than two individuals where the gender was all female and male, respectively. The ‘f+’ and ‘m+’ categories represent grouping of more than two individuals for which a female or male author were listed first in a mixed gender association. The plot shows a diminishing percentage of blue colored proportions indicative of a decreasing proportion of author associations with males in the first position.

For the 491 publications were three or more authors contributed equally, the most common form involved mixed gender contributions (348, 71%). Of these 200 listed a male author first while 148 listed a female author first. Comparing male first versus female first ratios observed versus expected values yielded p = 0.048. Although lower numbers precluded a decadal analysis these numbers imply a preference for male authors in the first positions of a multi-author byline when three or more individuals contribute equally. Analysis of the frequency of publications with three or more authors as a function of publication year revealed a positive trend line with time, suggesting that these author associations are becoming more frequent (Figure 2).

**Figure 2.**
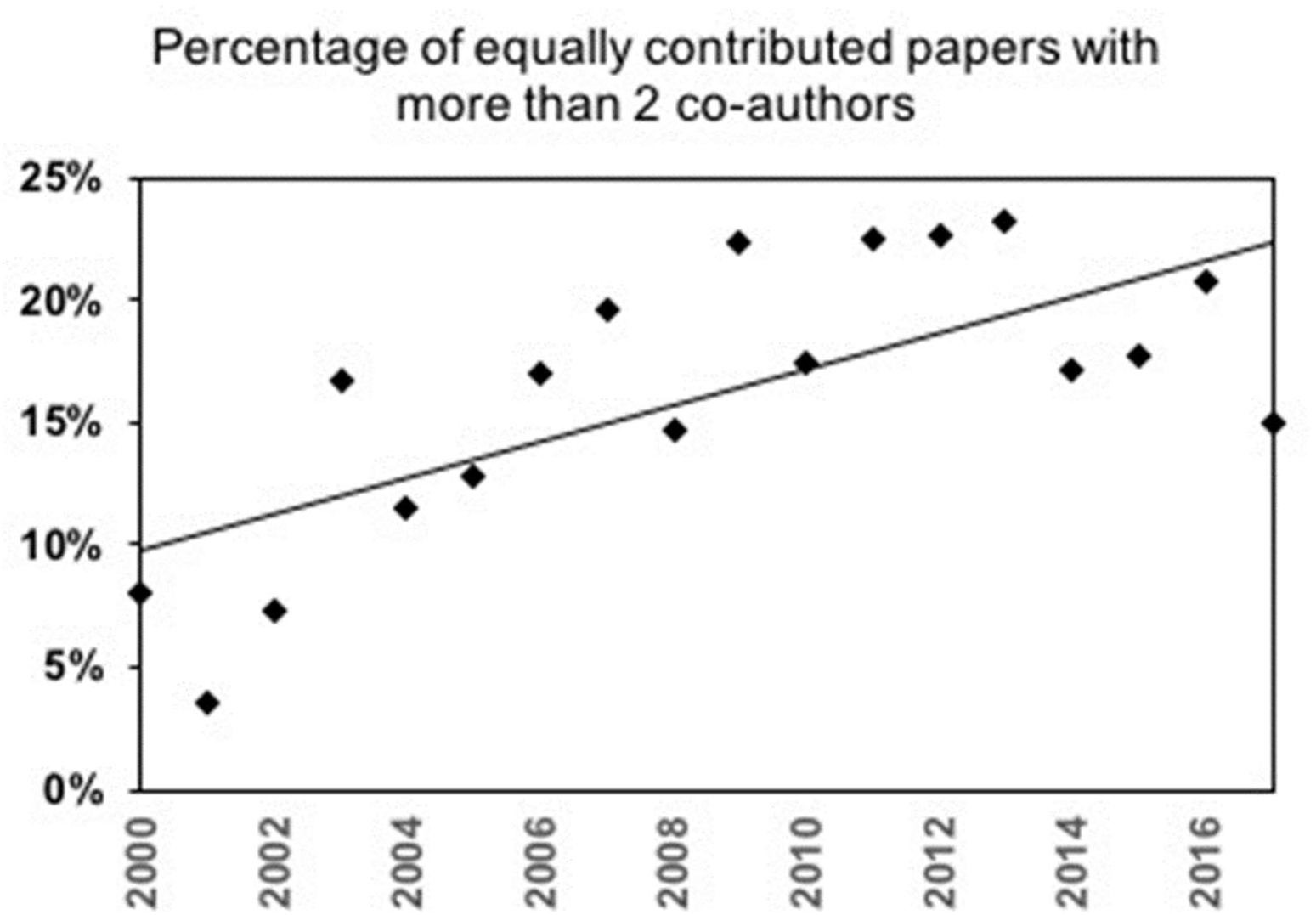
Percentage of paper with two or more individuals contributing equally as a function of time. Trendline R^2^ value was 0.4857.

Analysis of the relative distribution of the eight types of author association as a function of continent from which the publication originated revealed similar patterns for Asia, North America and Europe (Figure 3). Patterns for Africa, Europe, and South America were different from those of Asia, North American and Australia groupings, but some of these categories contain fewer papers, which suggests a need for caution in comparing between these continental groupings.

**Figure 3.**
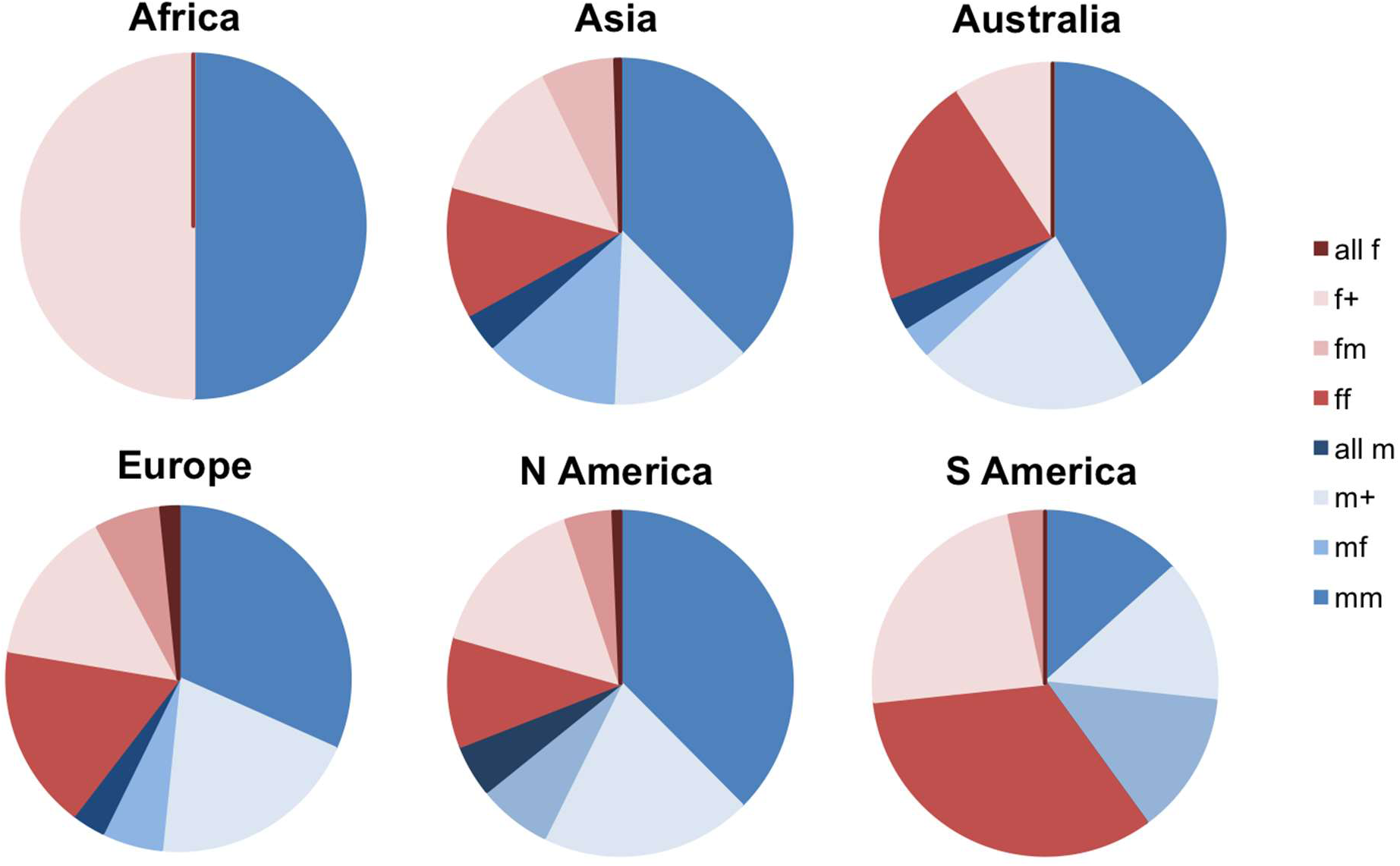
Pie diagrams indicating proportion of papers from six continents. The ‘fm’, ‘ff’, ‘mf’ and ‘mm’ categories represent pairings of female-male, female-female, male-female and male-male author orderings, respectively. The ‘all f’ and ‘all m’ represent groupings of more than two individuals where the gender was all female and male, respectively. The ‘f+’ and ‘m+’ categories represents grouping of more than two individuals for which a female and male authors were listed first in a mixed gender association. Papers were grouped based on country and continent association of corresponding author; Africa (n=4), Asia (n=221), Australia (n=65), Europe (n=988), North America (n=1523), and South America (n=30).

For 2109 of these author associations, we recorded author initials, which allowed to us analyze the frequency of alphabetical ordering for author sequences. The percentages of author associations for which the author sequence was alphabetical were 49%, 49%, 48%, 54%, 22%, 35%, 25% and 22%, for male-male, male-female, female-male, female-female, more than three authors all male, more than three authors all female, more than three authors mixed gender male lead or female lead, respectively. In comparing male vs female first position there was no significant difference between the frequencies of alphabetical versus no-alphabetical ordering.

## Discussion

Biomedical research is increasingly collaborative producing multi-authored publications [2]. Authorship is the major mechanism for crediting the contributions of individuals to a scientific study and offers of authorship are essential and expected when publishing the product of collaborative research. In addition to authorship, listing the position in the author byline of published scientific papers is an accepted form of credit allocation. The first position is reserved for the individual who has contributed most to a scientific publication. In recent years, the phenomenon of shared first authorship or equal contributions to the first author position has become increasingly common as a mechanism for sharing credit in studies where more than one individual makes critically important contributions to the study. In this study, we analyzed the author gender distribution and byline order for author combinations designated as contributing equally to a scientific publication. Given persistent inequities in the position of women in academia and that publication and authorship position are critical for employment, promotion and grand funding, it is important to know whether gender differences exist in the ordering of authors that contribute equally to publications.

Male-male associations were the most common gender combination and this applied to combinations involving both pairs and association of more than two authors. Female-female associations were the least common gender combination comprising less than half the number of observed male-male associations. Male-Female and Female-Male associations were almost as common as male-male associations, but the frequency differed in gender order. Male-female associations were significantly more frequent than female-male associations, with a ratio of 1.3:1. However, analysis of the data as a function of time revealed that the effect was strongest for publications dating before 2007. In the past decade, the preference for male gender in the first position of mixed-gender pairings has declined such that there was no statistical difference between observed gender order pairings and those expected from random assignments. Whereas pairings of two authors sharing equal contribution for the first author position comprised the majority of associations, we found a significant minority for associations involving three or more authors contributing equally. As with single pairings, the preference for male gender in the first position also occurred with author associations of three or more authors listed as contributing equally, with a ratio of 1.35:1. Male only associations were also more common when three or more authors contributed equally such that there were almost four times more male only associations than female only associations.

Although we have no information on the mechanism for selection of these gender author assignments the disequilibrium between observed and expected ratios strongly suggests that these selections were not made randomly. In fact, only one of the publications we analyzed provided the rationale for the author order and indicated it was based on alphabetical ordering (for example see [17]). Given the importance of the first author position in credit allocation for publications in the biomedical sciences the disparity in frequency between male-female and female-male associations raises the possibility for unequal gender benefit among associations sharing the first authorship despite these being designated as contributing equally. We note that information on equal contributions is often included as a footnote and that for some publications it is stated only in the print version. Consequently, it is likely to that the author listed as contributing equally in the second position may be not benefit as much as the author listed first, which is usually recognized as the most important contributor in biomedical publications.

The finding of a disproportionate number of males in the first position relative to expected numbers had these positions been selected randomly is consistent with several studies showing that females receive less credit recognition relative to their male colleagues. A study of PhD students revealed that male graduate students were 15% more likely to be listed in publications than their female counterparts [18]. An analysis of male and female authorship patterns for publications in natural sciences, social sciences and the humanities showed that a large predominance of male author over female authors in the first and last positions [19]. Perhaps most relevant for our findings is the observation that women receive less credit than men for team work in academia [20]. The finding that the preference for male first publications had declined in the past decade could reflect gains by women in academia in recent years. Nevertheless, given that authorship position in a scientific paper can have career altering consequences, choices made years ago could have long lasting effects that may still be a contributing factor to current gender inequities in academia.

We observed that male-only pairings predominated in author combinations of two or more authors. Again, without access to how these orderings were decided, or to the gender composition in the laboratories, we cannot infer the causes for this gender preference. Nevertheless this observation is consistent with the finding that males are more likely to share data with other men, which can lead to scientific discussions and collaborations that result in shared first author publications [21]. Similarly, the high prevalence of publications sharing first authorship among three or more males echoes the concern that male-exclusive networks exist in science [21].

The frequency of multi-author equal contributions dropped rapidly for associations of more than three authors but we observed at least two groupings of 11 authors [15; 16]. We noted a positive trend for the frequency of publications listing three or more authors contributing equally, suggesting that such author associations may be increasing as a function of increased team science in biomedical research. Although we have no information on the mechanism for sharing credit among so many authors, we note that some have questioned whether statements of equal contribution can ever be accurate given the problem of weighing the relative value of different contributions [17]. A recent analysis of journal instructions for authors revealed that none addressed equal contribution statements [18]. Our findings of a disequilibrium between observed and expected male and female authorship positions among groups of authors that contributed equally suggests a need for explicit requirements that explain how the ordering is done.

The majority of publications analyzed came from the United States, reflecting the predominance of this country in contributing to the biomedical literature. Analysis of distribution of author associations for the continent of origin produced similar patterns for Asia, Europe and North America, which may reflect similar practices in author order selection in the contributing countries. We note with interest that the patterns for Africa, Europe, and South America differed from the Asia, Australia, and North American groupings, but caution against drawing conclusions since for some of these continent groupings the number of publications analyzed may not be adequate to make direct comparisons. Nevertheless, the possibility that there are differences in author gender order and associations depending on country of publication is an interesting area for future investigation.

We acknowledge some limitations in our study, which suggest caution in interpreting the data. The finding that the preferences for males and females in the first author position varied over time, suggests that variables contributing to these decisions may be changing rapidly. We noted differences between journals in the proportion of pairings suggesting that that there may be differences between fields that could skew results depending on the source database. The approach to search citations can also affect the results depending on the method used. Using search engines such as Google Scholar facilitates searches since searching for the words ‘contributed equally’ in journal sites usually identifies many irrelevant publications where these words are in the text of the article. However, using the search engine introduces potential biases depending on how the algorithm prioritizes those publications containing the words ‘contributed equally’. Many of the results from those searches were not usable in this study because they related to shared internal and corresponding author contributions and to the use of the search phrase in the text of the paper. For some publications, authors were listed using first initials and it was not possible to assign gender. We could not reliably assign gender to authors from name alone in approximately 4% of publications. Given these limitations and caveats our findings should be considered preliminary until confirmed by subsequent studies, which may be able to analyze a larger number of publications across many disciplines through automated searches linked to gender image recognition software. Nevertheless, the finding that a disequilibrium exists in gender ratios among authors listed as contributing equally is sufficiently robust to raise concern on the fairness of shared credit contributions assignments in biomedical publications. At the very least, this study opens a window into a relatively unexplored area in the sociology of science that could have major consequences for current efforts to improve gender equity in academia.

In summary, our results provide evidence that the first positon of author bylines involving mixed gender associations contributing equally to a publication is more likely to have a male author. Given the importance of first authorship in biomedical publications and the increasing popularity of sharing authorship with the rise of team science a male preference could have consequences on hiring decisions, promotion and the distribution of resources such as grant funding. This information should be of interest to promotion and grant review committees as they consider the merit of applications who list papers stating that they contributed equally. The finding of disequilibrium in gender ratios among authors who contributed equally raises the possibility that some authorship decisions are vulnerable to conscious or unconscious biases and this suggests the need for journals to require statements of how author ordering was done in publications claiming equal contributions.

---

**Supplemental Figure 1.**
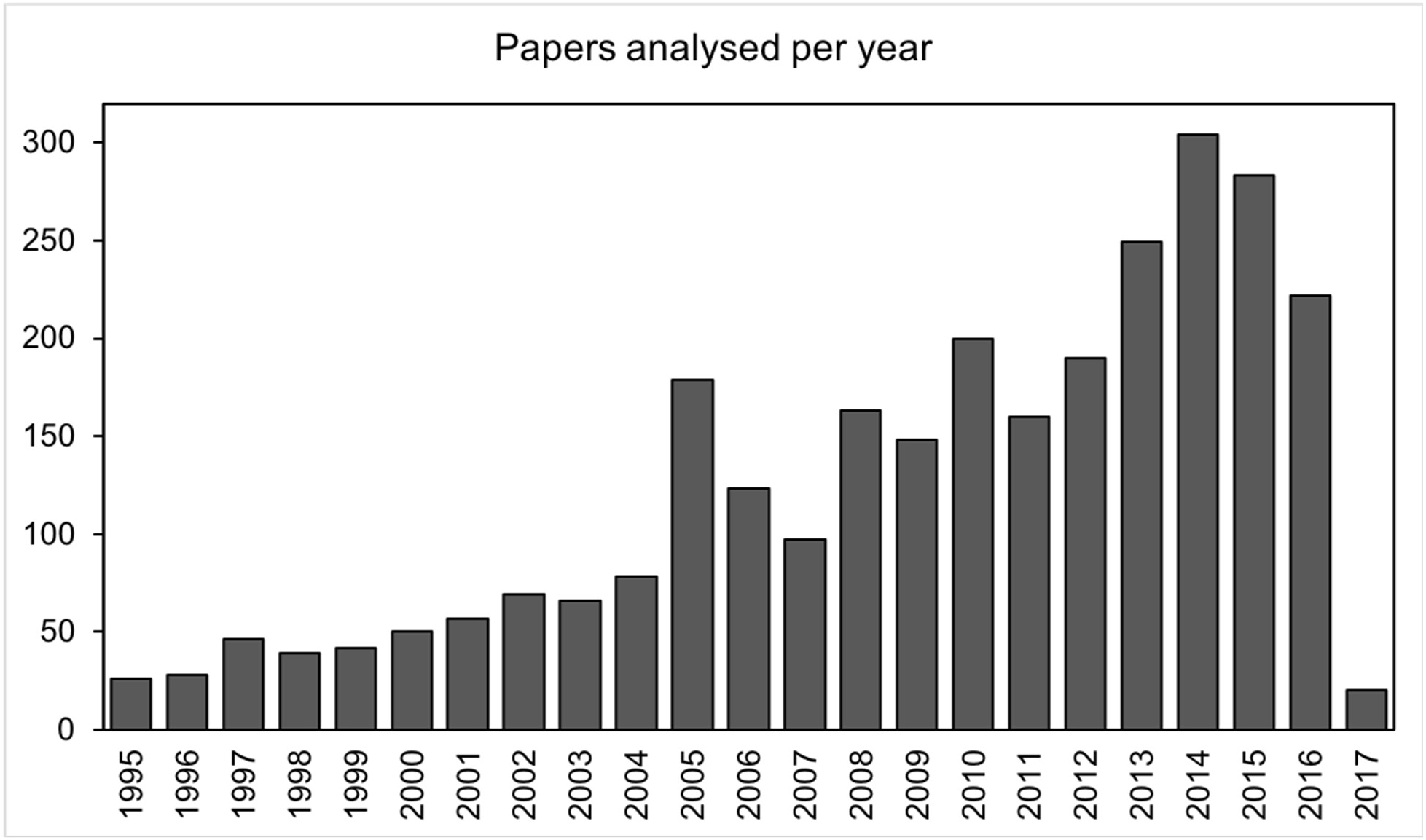
Papers analyzed per year in this study.

